# An ion channel omnimodel for standardized biophysical neuron modelling

**DOI:** 10.1101/2025.10.03.680368

**Authors:** Chaitanya Chintaluri, William Podlaski, Panos A. Bozelos, Pedro J Gonçalves, Jan-Matthis Lueckmann, Jakob H. Macke, Tim P. Vogels

**Affiliations:** Institute of Science and Technology Austria, Austria; Champalimaud Centre for the Unknown, Champalimaud Foundation, Portugal; VIB-Neuroelectronics Research Flanders (NERF), Belgium; Departments of Computer Science and Electrical Engineering, KU Leuven, Belgium; Google, Switzerland; Machine Learning in Science, University of Tübingen and Tübingen AI Center, Germany; Department Empirical Inference, Max Planck Institute for Intelligent Systems, Tübingen, Germany

**Keywords:** Voltage-gated ion channels, Compartmental models

## Abstract

Biophysical neuron modeling is an indispensable tool in neuroscience research, with the combination of diverse ion channel kinetics and morphologies being used to explain various single-neuron properties and responses. Despite this, there is no standard way of formulating ion channel models, making it challenging to relate models to one another and experimental data. Here, we revive the idea of a standard model for ion channels based on the Hodgkin-Huxley formulation, and apply it to a recently curated database of ion channel models. We demonstrate that this standard formulation, which we refer to as an omnimodel, accurately fits the majority of voltage-gated models in the database (over 3,000 models). It produces similar, if not identical, responses to voltage-clamp protocols in simulations where the ion channel omnimodels were used. Importantly, the standard formalism enables easy comparison of models based on parameter settings. It can also be used to make new observations about the space of ion channel kinetics found in neurons. Furthermore, it facilitates the inference of ion channel parameters from the responses to standard protocols. We provide an interactive platform to compare and select channel models, and encourage the community to use this standard formulation whenever possible, to facilitate understanding and comparison among models.

## 1 Introduction

Single neurons form the building blocks of neuronal signal processing, and the immense diversity of their activity profiles across brain areas and species contributes in myriad ways to the resulting circuit computations [1]. The electrical properties of single neurons largely stem from the composition and distribution of ion channels across the neuron’s morphology. They can lead to complex voltage dynamics and input-output functions [2, 12 3]. To understand their emergent properties, computational modelling has proven essential [4]. This, of course, dates back to the seminal work of Hodgkin and Huxley [5] in characterizing the voltage-dependent dynamics of sodium and potassium ions underlying action potential generation. Since then, the Hodgkin-Huxley (HH) formalism has been generalized to describe arbitrary voltage-dependent ion channel dynamics using sets of gating variables which can be qualitatively or quantitatively matched to data [6].

A considerable number of biophysical neuron models, including ion channel models, have been developed over the past several decades. They are stored online, in archives such as ModelDB [7]. With many channels of each type, it is very difficult to determine how these models relate to one another. To address this, we created a new database of ion channel models originally coded in the NEURON language [7], which features a more quantitative approach to comparing models (IonChannelGenealogy (ICG); [8]). ICG helps organize ion channel models according to various metadata such as animal species, neuron type, activating function, and publication pedigree. It also enables the comparison of the variability of ion channel models of the same kind. However, the programmatic implementation of the ion channel and its underlying biophysical variables varies in complex and unique ways, and there’s no clear or intuitive way to understand how the channels relate, or to choose the best one for a given application.

Standardization is a hallmark of open and collaborative science, and for computational modelling, it facilitates the understanding and reuse of models. Indeed, several previous studies have proposed the use of standard models for ion channels [6]. Despite this, no standard formalism has been adopted for ion channel models, and it is common to see a range of different functions and implementations thereof to express the steady-state activation and inactivation curves, as well as the time constants of these ion channel models. While such diversity enables flexibility in fitting experimental data, the lack of standardization makes it difficult to understand how the underlying ion channel parameters and dynamics give rise to a particular single-neuron phenotype, and how robust this phenotype is.

Here, we revisit the idea of a standardized formalism for ion channel dynamics, and follow Destexhe and Huguenard [6] to describe a standard model. We apply this formalism to over 3,000 models in the IonChannelGenealogy (ICG) database and find that we can emulate the majority of channel models with high fidelity, i.e., their behavior remains remarkably close to identical in voltage-clamp protocols and full neuron simulations. We refer to these new ion channel models, with standardized gating variables, as the corresponding ion channel “omnimodel”. We illustrate how the omnimodel formalism enables the comparison of parameters across different ion channel models, thereby providing a new dimension for ion-channel analysis. To facilitate translation to other models in the future, we provide specification sheets for each channel, including steady-state and time-constant curves, fitted parameters, and lineage information. We also traced how channel mechanisms have been reused and modified over the past 75 years by comparing their source code, revealing families of models that evolved through small edits or reimplementations. Finally, we built an interactive web application (https://icg-explorer.org/visualizer) that lets users explore these similarity networks in a force-directed layout.

## 2 Results

In the following, we fit a standardized thermodynamic ion channel model formulation based on the Hodgkin-Huxley model to a group of 3524 ion channel models on the ICG database (originally from ModelDB). All models are voltage-gated and can be categorized into five classes based on ion channel species: potassium (K, 1455 channels), sodium (Na, 923 channels), calcium (Ca, 633 channels), nonspecific cation (Ih, 250 channels), and calcium-dependent potassium channel (KCa, 263 channels).

### 2.1 Extracting steady states and time constants data points from .mod files

Ion channel models in the HH formalism are composed of multiple gates. From each ion channel .mod file, we extracted 1) the names of the gates, 2) the power of each gate, 3) each gate’s steady state activating (or inactivating) curve, which captures the probability of it being engaged at a given steady-state voltage, and 4) its time constant curve – the voltage-dependent time constant for each gate. The gating variables (3, 4) were extracted at discrete physiological membrane potential values (between −100 *mV* and 100 *mV*) and at three different temperatures (6.3^*°*^*C*, 23^*°*^*C*, and 37^*°*^*C*). In the case of calcium-dependent potassium channels, we extracted all variables at three intracellular Ca^2+^ concentrations (0.05*µ*M, 0.5*µ*M, and 5*µ*M, See Methods). In all, we successfully extracted the steady states from 1266 K channels (1629 gates), 808 Na channels (1338 gates), 567 Ca channels (806 gates), 211 Ih channels (170 gates), and 231 KCa channels (247 gates), with a total of 3083 channels (87% of all available channels on ICG.) and their 4190 gating variables (Figure 1).

**Figure 1:**
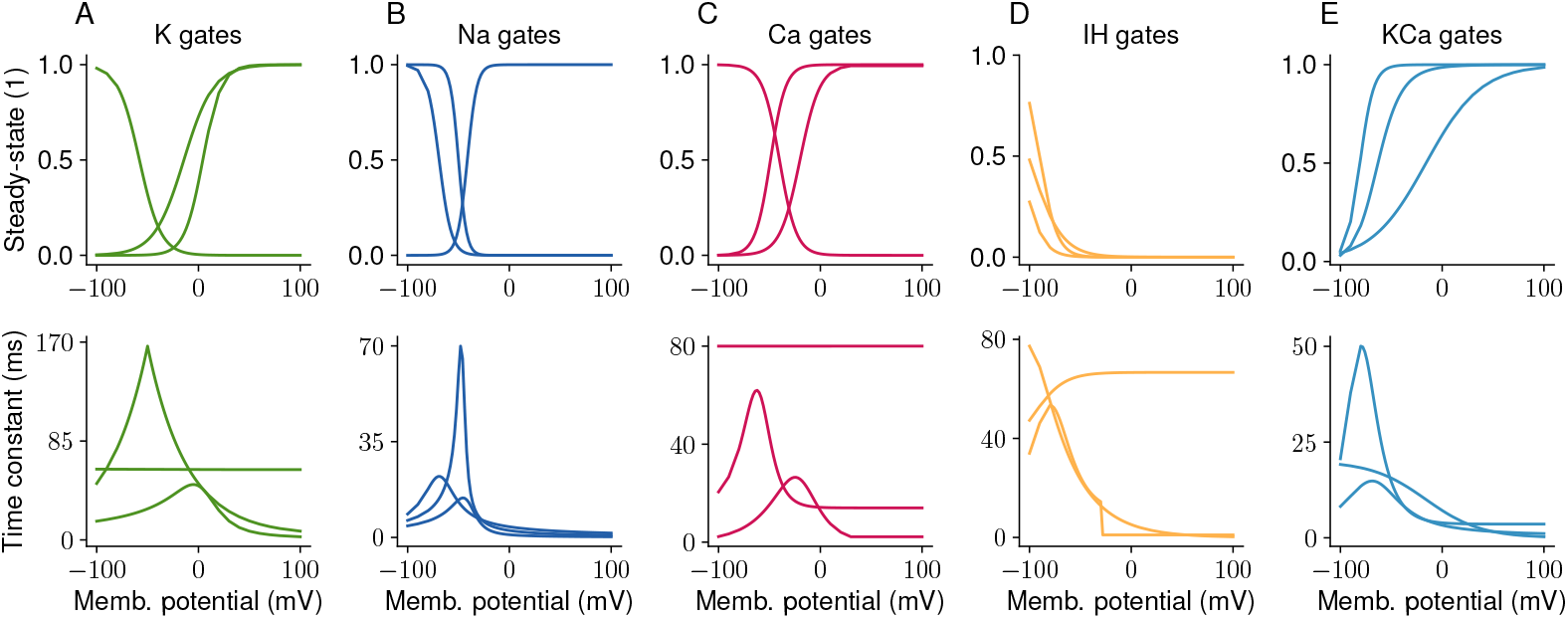
A sample of three gating variables for the sub-families of voltage-gated **A)** potassium, **B)** sodium, **C)** calcium, **D)** Ih- and **E)** Ca-dependent potassium channel models available on ICG. These gating variables were extracted directly from the respective .mod files at 6.3^*°*^*C* and 5*µ*M intracellular Ca^2+^ concentration. The top row shows the probability of the channel opening at different membrane potentials, and the bottom row shows the corresponding time constants at these potentials.

### 2.2 Omnimodel: a standardized formulation of ion channels

The standardized formalism for ion channels, developed by Destexhe, A., and Huguenard, J. [6], expresses two components for each gating variable: a steady-state activation (or inactivation) curve, and its corresponding time constant curve. Following this, we describe the steady-state activation (or inactivation) curve as a sigmoid with two parameters (*a,b*):

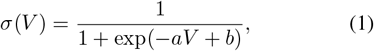

where *a* determines the slope of the curve, and *b* the membrane potential at 0.5, often called the V-half value. To increase the flexibility of the model, we also tested a modified sigmoid curve with two additional parameters (*c, d*):

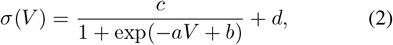

where *c* sets a multiplicative correction of the curve, and *d* determines its offset.

The time constant curves in the population of ion channel models are more varied (Figure 1, bottom row), ranging from sigmoidal to non-monotonic single peak, and even more complex shapes. To capture this diversity of time constant functions, we formulated a time-constant model with ten parameters (*vh, A, b*1, *c*1, *d*1, *b*2, *c*2, *d*2, *e*1, *e*2),

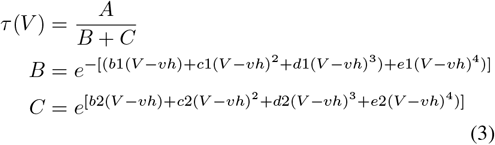

which is a fourth-order thermodynamic formulation [9]. To assess the sufficiency of the formalism, we incrementally increased the order of the terms in the denominator, starting from a first-order denominator (i.e., *c*1, *d*1, *e*1, *c*2, *d*2, *e*2 set to 0) up to the fourth order. We refer to the composition of these standardized gating variables into an ion channel model as an *omnimodel* of that particular ion channel.

### 2.3 Fitting standardized curves to data

The gating variables, i.e., the voltage-dependent functions of steady state and time constants, were obtained at discrete membrane potentials directly from the NEURON .mod files. With this data, we could now fit a standardized formulation for each omnimodel gate and obtain the coefficients of the desired functional forms in an automated way, regardless of their original formulations in the .mod files. We used non-linear least squares to fit the data extracted for each gate of the ion channels. This approach was successful for approximately 72% (success/total, K:1016/1266, Na:546/808, Ca:429/567, Ih:167/211, KCa: 71/231) of models. This amounted to 2229 ion channel omnimodels in all. The other ion channels could not be fit or were excluded due to run-time errors when fitting the curve to our formulation for at least one of the gates (Figure 2).

**Figure 2:**
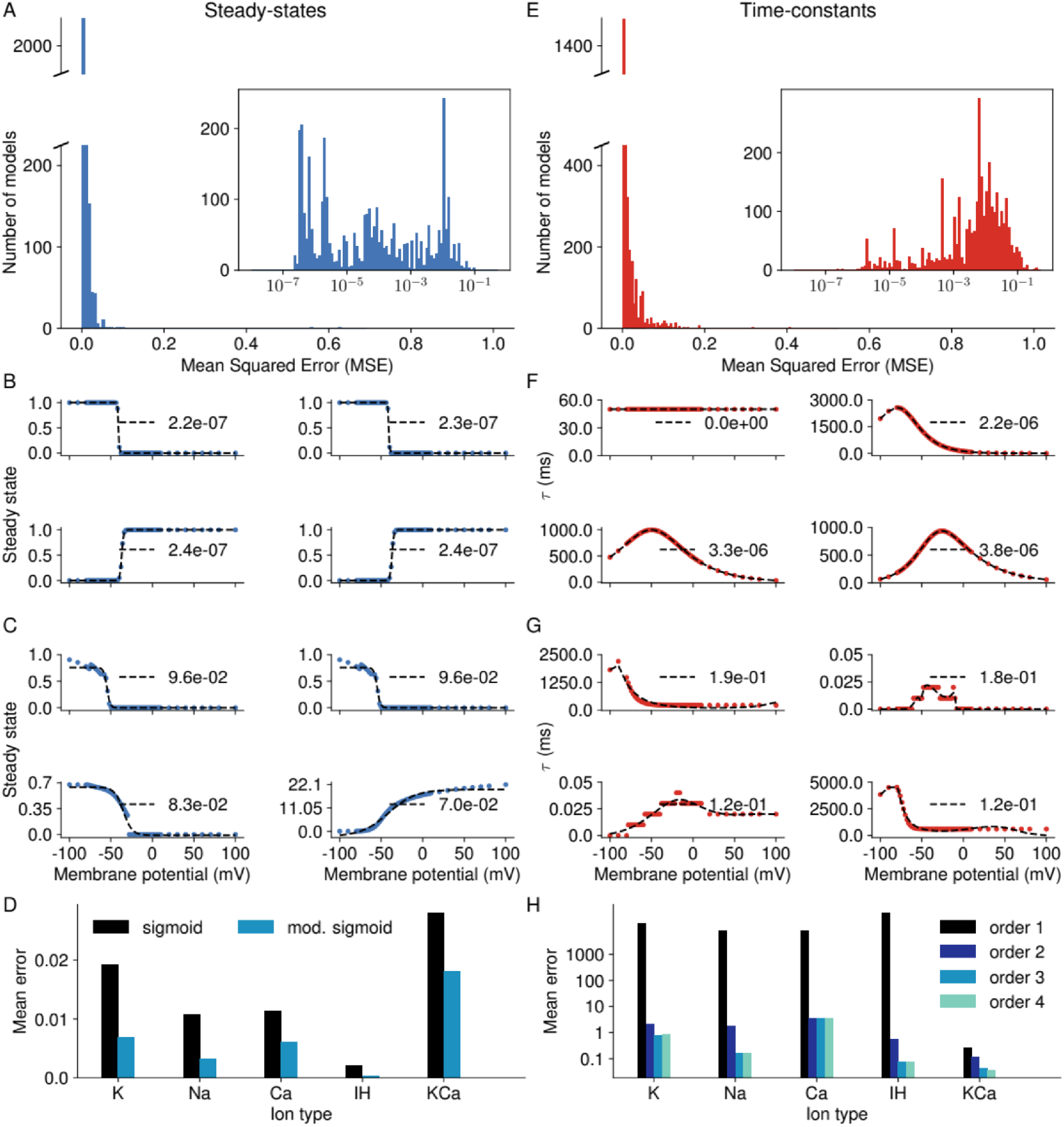
Summary of the manual fitting procedure for all the channel variables. **A)** Histogram of the errors between the modified sigmoid and the original activation function of all channel models that produced a fit at all. (Eq. 2). **B, C)** Comparison of numerically extracted gating values from the channel .mod files (dotted lines) and the corresponding fit (dashed line). **B)** Example fits with low errors, i.e., the steady state curve fits are good. **C)** Example fits where the error is relatively large, i.e., the fit is poor. **D)** The summary of the mean errors of all the model fits whose steady states were extracted from ModelDB. Here, sig is the sigmoidal formulation and m-sig is the modified sigmoidal fit. **E)** Histogram of the errors between the 3rd order fit (Eq. 3) and the original time constant function extracted from .mod files., followed by good **(F)** and bad **(G)** fit examples. **H)** The summary of the mean errors of the model fits for the time constants that were extracted from ModelDB.

The steady-state curves of the ion channel gates were fitted with a sigmoidal shape and correspond to either activation or inactivation gating properties. For most gates, the sigmoid fit was sufficient (Figure 2A, B, D). Bad fits were almost exclusively due to idiosyncratic formulations of the models, and as such, these models did not fully activate or inactivate, or in some cases, were non-monotonic curves with peaks (2C bottom). Similarly, the fits to the time constant curves (although with a slightly higher error than the steady-state curves, Figure 2E, H) fit most of the gating variables. Good fits (Figure 2F) were bell-curved in shape, and bad fits (Figure 2G) were usually due to very sharp, at times non-continuous changes in the time constant curves, which were difficult to fit with a continuous function.

Although our approach was generally successful in fitting the diverse range of ion channel models, it was unclear how small fitting errors would affect the performance of an omnimodel in practice, i.e., when combined into an interacting set of channels within a neuron model. We thus decided to evaluate the performance of the omnimodels in more realistic protocols and experiments.

### 2.4 Comparing omnimodels to original models using voltage-clamp protocol output

The first step in evaluating the performance of the models in biologically realistic simulations was to test their behavior in response to a series of voltage-clamp protocols, as previously designed for the ICGenealogy database [8] (Figure 3A). We employed five protocols - Activation, Inactivation, Deactivation, Ramp, and Action Potential- to thoroughly investigate the dynamics of the channel models in physiologically realistic voltage ranges. To directly compare the original and our omnimodels, they were simulated separately in NEURON, normalized to a max current of 1, and compared.

**Figure 3:**
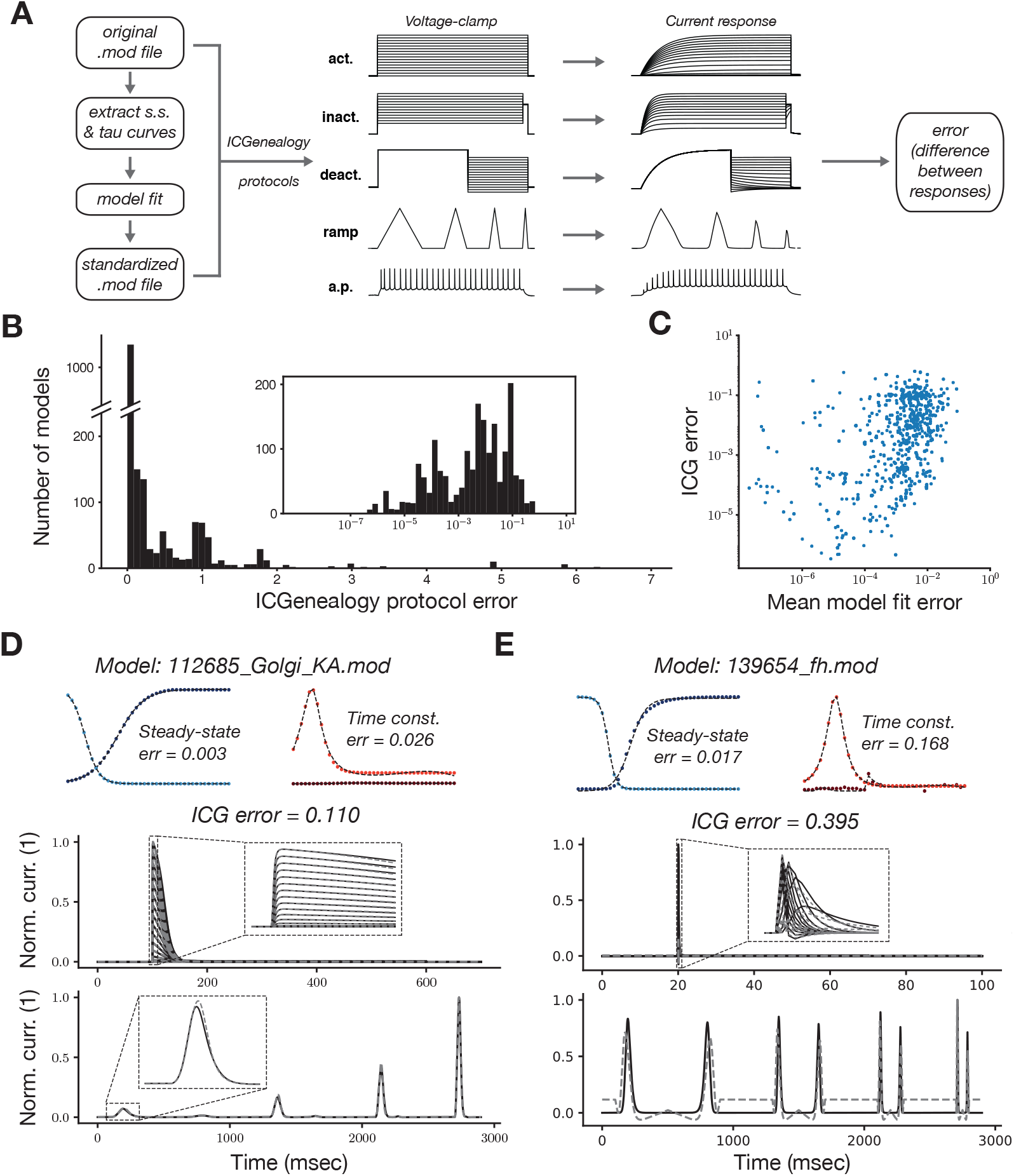
Deviations between the responses of the omnimodels and the original models for established stimuli. **A)** Schematic of the procedure of extracting the steady state and time constant functions from original models and recomposing them into a new .mod file, and comparing the two models by way of the voltage clamp protocols used in ICG. **B)** A histogram of the deviations in the ICG protocol response between the original model and the omnimodel. Inset showing logarithmic scale. **C)** Correlation between the fitting errors of the gating variables (Figure 2) and the ICG protocol response deviations. Worst case examples: **D)** Sample ion channel with good fit of the gating variables (top) but large response deviation (bottom) for activation and ramp stimulus between original (solid) and the omnimodel (dashed) **E)** Sample ion channel with poor fit of the gating variables (top) and large response deviation (bottom) for activation and ramp stimulus between original (solid) and the omnimodel (dashed).

We observed that most omnimodels were indistinguishable in their current response from that of their .mod file counterpart (Figure 3B, D). For the less well-matched models, the magnitude of the mismatch between original and omnimodel was correlated with the fitting errors for the steady state and time constant curves (Figure 3C). Mismatch in performance could occur, e.g., in cases where the sigmoid activation curve of a channel was not truly sigmoidal Figure 3E), or when the original time constant function had slightly different slopes for the upswing of the curve than for the downswing. Given the success of these formulations, we then proceeded to test the performance of multiple omnimodel ion channels in previously published single neuron models.

### 2.5 Performance of combinations of omnimodels in single neuron models

We evaluated two single-neuron models from a selection of previously published, biologically plausible neuron models in ModelDB by replacing, when possible, the active ion channels in those single-neuron models with their corresponding omnimodels.

Towards that goal, we used a layer 5 pyramidal neuron model [10] that produces backfiring action potentials (Figure 4, top row). The original model contained 10 ion channel models (Ih, Im, NaT, NaP, CaLVA, CaHVA, KT, KP, SK3.1, SKE2), all of which we replaced with their corresponding omnimodels, except for two channels (SKE2, CaHVA). We tested the neuron model’s response to three published protocols (Figure 4 bottom row). First, we used a current injection into the soma, followed by a current injection into the dendrites 5ms later. This protocol resulted in a somatic voltage response (black), as well as a proximal (orange) and a distal (red) dendritic response (Figure 4A, same as Figure 4A in the [10]).

**Figure 4:**
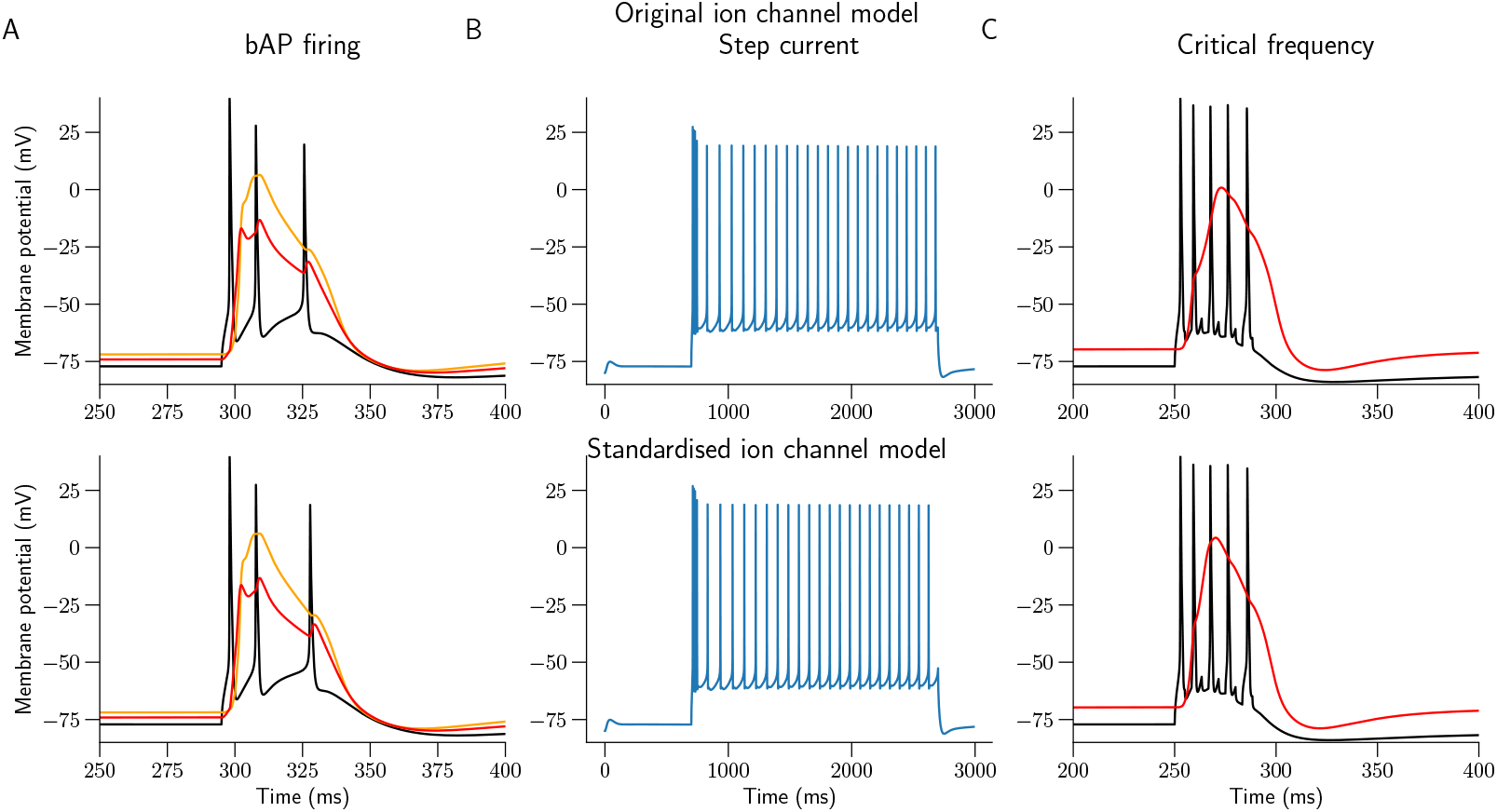
Indistinguishable responses from a layer 5 pyramidal cell model with ion channel models with idiosyncratic (top row) and omnimodel (bottom row) formulation, experiencing back propagating action potentials (bAP). **A)** A current injection protocol that is a combination of somatic and dendritic currents separated by an interval of 5 ms, evoked a proximal (orange) and distal (red) dendritic Ca2+ spike, followed by a burst of two additional somatic APs (black). **B)** The models’ firing response to a 793 pA depolarizing somatic step current for 2 seconds. **C)** High frequency stimulation (120 Hz, black) evoked a regenerative Ca2+ spike in the dendrites, as seen by membrane potentials from apical dendrites (red trace) in both cells.

Responses were indistinguishable between the original model (Figure 4 top row) and the omnimodel (Figure 4 bottom row). Second, we used a two-second-long depolarizing current (Figure 4B in the [10] paper) that again evoked near-identical spiking responses from both models (Figure 4 B). Finally, a high frequency stimulation of 120Hz resulted in a Ca^2+^ spike in the dendrites, also near-identical in the two models (Figure 4 C and Figure 5A in the [10]).

**Figure 5:**
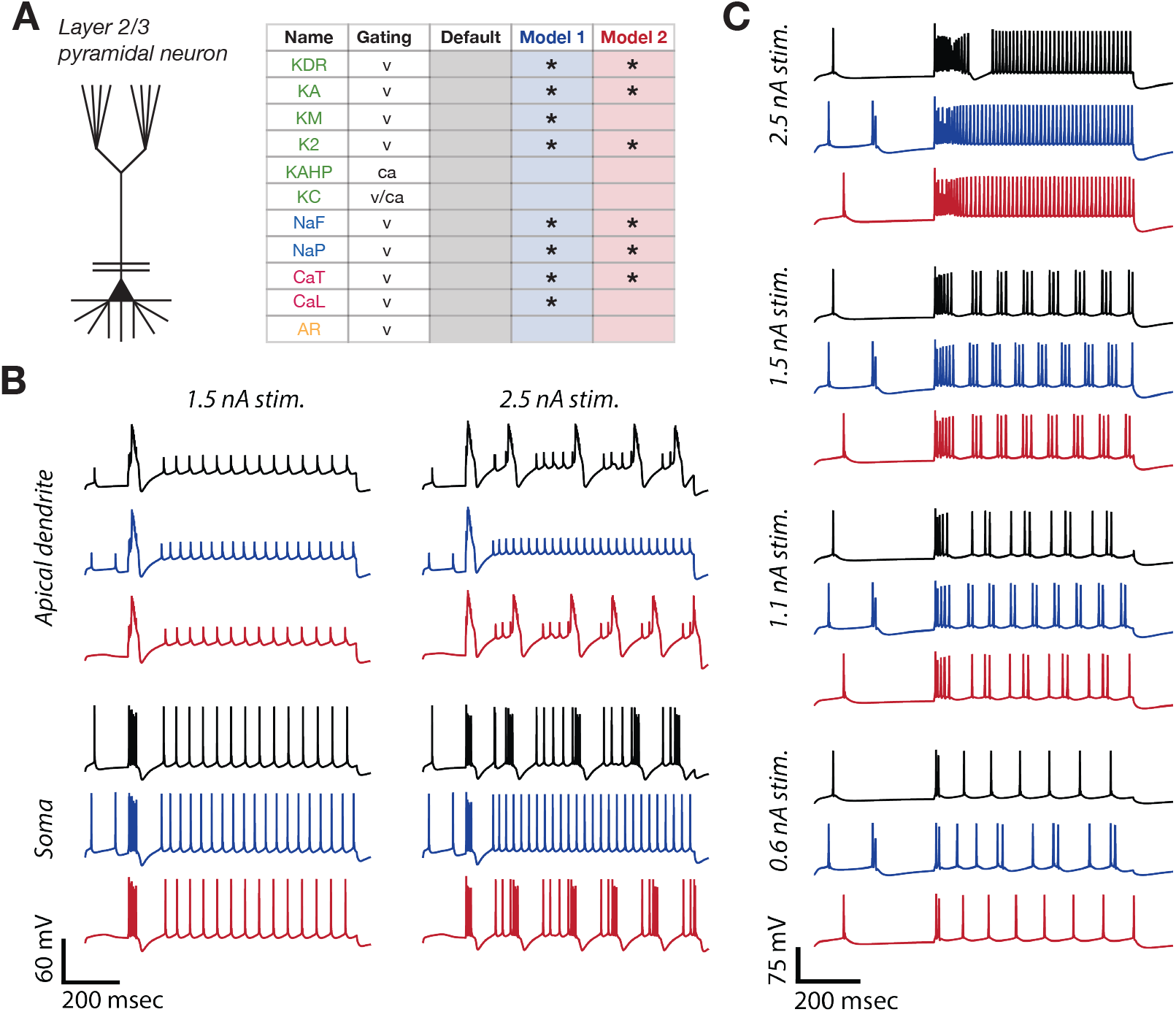
Comparison between a layer 2/3 pyramidal neuron model with original ion channels and several omnimodel replacements. **A)** The morphology of the neuron model, and a table of included ion channels with their gating variables (v = voltage, ca = calcium, v/ca = voltage and calcium). Stars in the fourth and fifth columns indicate which channel models were replaced with omnimodels in Model 1 (blue) and Model 2 (red). **B)** Membrane potential responses in the apical dendrite (top) and the soma (bottom) to a current stimulus of 1.5nA (left) and 2.5nA (right). **C)** Rhythmic bursting followed by fast tonic firing in response to various somatic current depolarization stimuli.

The second model we simulated was a layer 2/3 pyramidal neuron [11] in which we tested the response of the model neurons to depolarizing current pulses and obtained the resulting membrane potential in the soma and the apical dendrite (Figure 2 in [11]). We also tested the appearance of rhythmic bursting followed by fast tonic firing with increasing somatic depolarization (Figure 4 in [11]).

Here, we expected some complications because three of the omnimodel channels could not be fitted to the original activation functions. Additionally, two more of the omnimodels were relatively poor fits (KM and CaL). We thus compared the original neuron model with two variants of omnimodel combinations. In one model (Model 1, blue-colored traces in Figure 5), we replaced all but three ion channels (KAHP, KC, and AR) with their corresponding omnimodels. In the second model (Model 2, red-colored traces in Figure 5), we retained two other original channels (KM and CaL). Both models 1 and 2 were able to reproduce the results of the original model quantitatively with substituted omnimodels.

However, we noticed differences in some aspects of the spike waveforms, specifically in Model 1, which was unable to reproduce burst-like activity in the apical dendrites for the 2.5nA stimulus, and spiked at a higher frequency compared to the original (Figure 5 blue traces, B top-right, C top). Model 2, which incorporated CAL and KM channels from the original model, fared substantially better (Figure 5 red traces). As many neuron models feature some amount of parameter tuning using additional degrees of freedom (e.g., the spatial distribution of ion channel conductances), it is unclear to what extent these discrepancies reflect essential differences in dynamics or if they could instead be remedied through a more straightforward re-fitting procedure (see Discussion).

### 2.6 Comparison of omnimodel coefficients

Having fitted the standardized models and validated that they exhibit nearly identical behavior to the original models, we next examined how our new model formulation facilitated the comparison and evaluation of ion channel models in the database. Since all models now had a standard formulation, we could explore the parameter values across different metadata items to identify trends. However, we noticed that the omnimodel coefficients for the extracted gating variables formed tight clusters. Further analysis revealed that these clusters consisted of ion channel models that were either reused and renamed or minimally edited, nearly identical channel mechanisms.

To test this systematically, we retrieved all the ion channels, irrespective of their type, that were available on ModelDB (9,642 .mod NMODL files) and compiled a graph representation of 3,920 unique source-code nodes linked by 28,857 similarity edges (threshold *≥* 75%). We identified 5,722 redundant instances (59.3% of files), i.e., exact copies, corresponding to an average reuse factor of 2.46*×* per unique implementation.

To provide a means for inspecting and visualizing the provenance of an ion channel, we developed a website that allows users to view all the ion channels we have processed (Figure 6). The website will enable users to select ion channels based on their type (K^+^, Na^+^, Ca^2+^, KCa, Ih, or all) (Figure 6) A), where the area of each node corresponds to the number of clones (exact copies). Refinements in the selection could be made based on whether the models have an omnimodel and by choosing the code string similarity percentage to be considered as the nearly identical copies of the nodes. This filtered view is shown in (Figure 6B). Additionally, the gating variables of the selected node’s model, available on the channel’s GitHub page, can also be navigated to directly from this webpage (Figure 6C, D). Furthermore, details of the node currently selected can be displayed, which include all the extracted information about the channel node, including its clones (Figure 6E). Lastly, Multi-node selection can be made using *Rectangle select* (click and drag to draw a rectangle) or by clicking on nodes while holding the Shift/Command key. The specific changes between any two (or more) nodes can be shown in a pop-up window (after pressing the d key).

**Figure 6:**
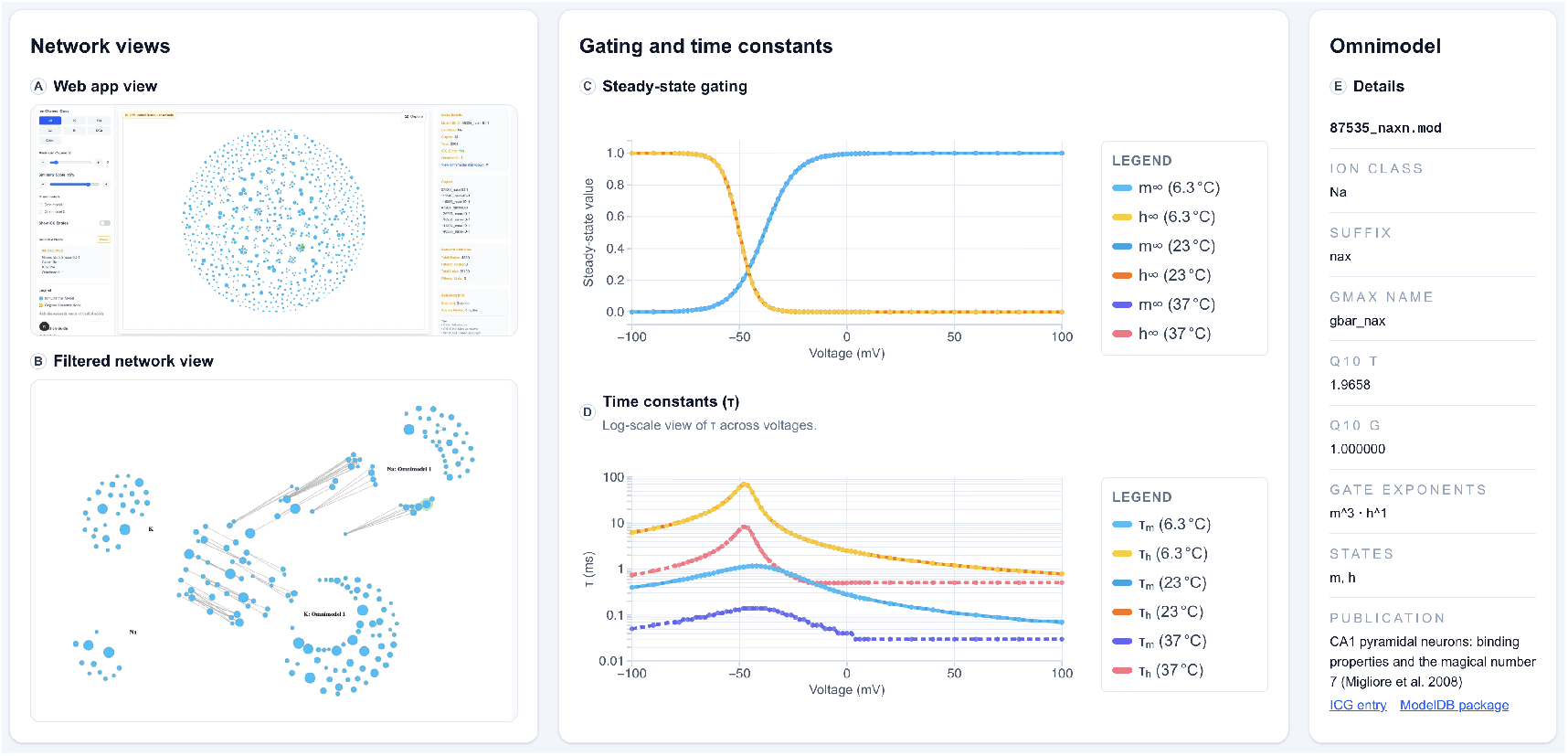
Interactive source–code similarity network and omnimodel metadata. **A)** Web app overview. **B)** Filtered network with class/ICG/omnimodel filters (K+ and Na+ classes are on and further grouped by omnimodel availability). **C)** Example of steady–state gating curves for the sodium channel file 87535_naxn.mod: *m∞* and *h∞* at 23°C (solid), with 6.3°C and 37°C variants (dashed); inline legend sorted by temperature (6.3, 23, 37°C). **D)** Corresponding time–constant curves *τ* (*V*) from `87535_naxn.mod` (log–scale *y*–axis). **E)** Omnimodel details: ion class, NMODL suffix, *g*_max_ symbol, gate exponents/states, and *Q*_10_ factors for *τ* and *g*; direct links to the corresponding ICG entry and ModelDB package.

### 2.7 Ion channel specification sheets

To make all the extracted information on ion channels reusable outside the web app, the ICG website now lists per-channel specification sheets (as an omnimodel.md file). Each sheet records: (i) ModelDB and ICG identifiers, (ii) ion class and NMODL suffix, (iii) gate set (names, exponents, and state count), (iv) steady-state *x∞*(*V*) and (v) time-constant *τ* (*V*) curves sampled at 6.3°C, 23°C, and 37°C, (vi) *g*_max_ measured at 6.3°C and 37°C together with *Q*_10_ factors for *τ* and *g* when available, (vii) omnimodel parameters with fitting errors (steady states and time constants), and (viii) source–code clone membership: pool ID, canonical “original” within the pool, and the full list of identical copies with links back to their ModelDB locations. These sheets are mirrored in the `icgenealogy` GitHub repository and referenced in the channel’s GitHub repository folder. Aggregated summaries, including gating variables and omnimodel coefficients, are also provided as Python pickles for programmatic use. The net effect is practical de-duplication: modelers can select one canonical implementation per clone family and avoid re-running analyses on files that are, in effect, the same channel.

## 3 Discussion

### 3.1 This paper and its applications

Building on the ion channel genealogy [8], we inspected individual ion channels and the underlying composition of the different gates, i.e., the number of gates that compose into an ion channel. When possible (for Na, K, Ca, Ih, and KCa ion channel families), we extracted the steady-state and time constant curves for each gate. We sampled these values at three different temperatures. We also obtained the maximum conductance values and the corresponding reversal potentials.

From these steady-state curves, we fit standardized gating variables (based on [6], [9]), i.e. *omnimodels*, whose parameters should sufficiently describe the gating variables and hence the ion channel behaviour. To test the accuracy of the omnimodels, we ran voltage clamp protocols to obtain activation, inactivation, deactivation, ramp, and action potential response curves, and compared their responses to those of the original model. Furthermore, we tested these models in two well-established single-neuron models by comparing the corresponding ion channels’ omnimodel outputs to the original. We found that the omnimodels effectively substitute for previous ion channel models.

While the goal of this effort was to extract gating variables from existing ion channel models, we note that the standardization provides a set of parameters that allowed us to compare all existing ion channel models systematically. Furthermore, this formalism has already enabled the development of an inference-based model [12], making it substantially easier to create new models that directly capture observations from experiments.

To document the lineage of existing ion channel models, we also performed a one-to-one comparison of the source code for all available ion channels. We observed that a substantial number of ion channels were reproductions of each other, as-is or minimally rejigged from some proto-ion channel models in the literature.

Such an approach may raise concern if these ion channels, initially constructed to fit particular existing experimental data, are modified or fine-tuned without specific ion channel information to justify these changes, without indicating such aberrations. We thus developed tools to inspect the similarity between ion channel models and compare their differences. We envision that this would enable new model developers to pick relevant ion channels for their study while also being aware of their history and the history of their modifications.

### 3.2 Future of ion channel modeling

Our work retraces a rather byzantine course of ion channel modeling in the past which did not follow a standardized convention when describing the channel properties, and relied on a somewhat anarchic community.

Many published models lack accompanying metadata, exhibit inconsistent temperature dependencies, and have other limitations, including arbitrary discontinuities in their formulation and unexplained changes to *vh* (Eq.3). In some cases, ion channels were presented as remixed versions of existing models, with minimal accompanying justification in the Methods. Some recent monumental efforts (notably NeuroML [13–15]) have made significant contributions to address these issues. Despite this, a large portion of existing ion channel models remains to be translated into these new formats. Our work provides the means to translate these old models into newer modern formats functionally ([16, 19 17]) and investigate their applicability to more contemporary data.

We emphasize that this study was limited to voltage-gated ion channels in the HH formalism on ModelDB [7]. While this represents a large dataset of ion channel descriptions, it is still dramatically incomplete compared to all ion channels expressed in the central nervous system and modeled in previously published work. For instance, even within ModelDB, we did not include hidden Markov models [18, 21 19] or other types of computational models [20–24]. Furthermore, we also excluded any metabotropic ion channels from our study [25, 29 26] as well as ion channels expressed in other organelles [27, 27 28] and cell types.

When examining biological realism, we recognize that expressed ion channels are not necessarily homomeric - i.e., composed of the same *α* sub-units, but are heteromers expressed as a re-combination of different *α* sub-units [29]. Other variants, depending on the subunit interactions, could change the gating properties of the ion channel [30]. The situation is even more nuanced when considering the *β* sub-units of ion channels, which can also modify channel dynamics [31, 32]. In recent years, it has become apparent that ion channels, in addition to voltage gating, may be sensitive to many other cellular variables such as redox, membrane-bound phospholipids, pH, intracellular ATP levels, etc [33–35].

We may think of the ion channels expressed in neurons as a meadow of wildflowers, each with multicolored petals that open only for a particular insect flying under moonlit skies. Yet, as a scientific community, we have so far glimpsed them only in muted tones, stripped of their rich cellular interdependencies. In this work, we aim to cast these channels in a more multispectral light. To that end, we have systematically cataloged a small subset of ion channel models assembled by those before us. We hope that future generations of ion channel enthusiasts will use this catalog to recognize and address the limitations of our current approaches, and in doing so, build more effective frameworks to explore the intricate meadow of ion channels that still lies before us—beautiful, complex, and mysterious.

## 4 Methods and Materials

### 4.1 Hodgkin and Huxley conduction-based ion channel models

In Hodgkin-Huxley models, also called conductance-based models, the ionic current passing through a channel is proportional to the difference between the membrane potential and the reversal potential of that ion. Additionally, the current is proportional to voltage-dependent ‘gates’ which are either activating (allowing ionic flow) or inactivating (disallowing ionic flow) when engaged.

For instance, in the original Hodgkin-Huxley formulation for the Potassium channels, the total Potassium current exiting the membrane through these types of channels was given by

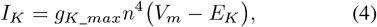

where *g*_*K*_*max*_ is the maximum conductance, *n* is the voltage-dependent activation gate which controls the current, whose value ranges between 0 when disengaged and 1 when engaged, *V*_*m*_ is the membrane potential, and *E*_*K*_ is the reversal potential of potassium ions. Experimental data can be best fit using four activating gates, hence the power of 4. Similarly, the sodium channel’s response could be fit using three activating gates (m) (hence the power of 3) and one inactivating gate (h), and their corresponding maximum conductance *g*_*Na*_*max*_, and reversal potential for sodium ions being *E*_*Na*_. The net current exiting the membrane was given by

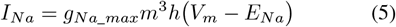

The success in expressing these currents to explain the spike shape in a squid giant axon prompted several other ion channel types to be formulated using similar gates. For instance, the fast activating and inactivating K type channels (also called K A-type) include an inactivating gate. The common aspect among all these gates is that they are voltage-dependent and could be expressed generally as follows

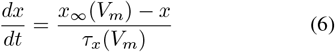

where *x* = *{n, m, h, etc*., *}*, and *x*_*∞*_ (*V*_*m*_) is the voltage-dependent steady-state value towards which *x* evolves with a voltage-dependent time constant *τ*_*x*_(*V*_*m*_).

### 4.2 Extraction of steady-states and time constant curves from .mod files

We use the script *HHAnalyse* from the pyNeuroML library [15] to extract the steady-state and time constant curves from HH channel models using the Neuron simulator. Each ion channel is inserted into a neural segment of length 10 *µm*, and diameter 5 *µm*, with passive conductivity of 0.001 *µS/cm*^2^ and reversal potential of -65 *mV*. After setting a simulation temperature of either 6.3^*°*^*C*, 23^*°*^*C*, or 37^*°*^*C*, we ran a voltage clamp protocol (holding voltage between −100*mV* and 100*mV*, in steps). We obtained all the gating variables (*x*’s) as they settled to their respective steady-state values. Once the slope of these gating variables approaches zero, we record the steady-state value and the total time taken to reach this value at that holding voltage. From this, for each gate in the ion channel we obtain their voltage dependent steady-state - *x∞* (*V*_*m*_) and their temperature and voltage dependent time constant curves *τ*_*x*_(*V*_*m*_, 6.3^*°*^*C*), *τ*_*x*_(*V*_*m*_, 23^*°*^*C*) and *τ*_*m*_(*V*_*m*_, 37^*°*^*C*).

### 4.3 Extracting maximum conductance from .mod files

We observed that the ICG protocols (see Section: Running ICG Protocols) we used to evaluate the robustness of ion channels depended strongly on the maximum conductance at which we were testing. To address this, we extracted the precise maximum conductance assigned to the ion channels in the original models using the Neuron simulator. For this, we insert the ion channel of interest into a neural segment of length and diameter of 5.64 *µm* (whose effective surface area was 100 *µm*^2^). The simulation temperature was set to either 6.3^*°*^*C* or 37^*°*^*C*, and we simulated without any additional stimulation. We note the net current exiting the membrane at the last time point.

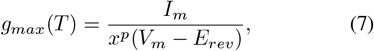

where *g*_*max*_(*T*) is the maximum conductance at temperature T (see next section), *I*_*m*_ is the trans-membrane current through the membrane, *V*_*m*_ is the membrane potential, and *E*_*rev*_ is the reversal potential of the ion, which we set. *x* here represents the steady-state value of the gate, and *p* is the number of these gates. At the end of the simulation, despite the temperature difference, the variables *x* and *p* would remain unchanged.

### 4.4 Temperature dependence

In the HH models described in ModelDB and our database, 51% (1587 of 3083 models) of all the models we tested include some temperature dependence. In these models, the dependence has been formulated in two different ways. Firstly, a temperature-dependent correcting factor (*t*_*adj*_) is used to adjust the channel density[36]. This conduction *fudge* factor effectively ensures that at higher temperatures, a greater number of ion channels participate in the membrane dynamics. This is modelled as a multiplying factor to the maximum conductance, i.e., *g*_*max*_(*T*) = *t*_*adj*_ *\* g*_*max*_, where *t*_*adj*_ is given by:

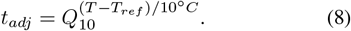

Here, *T*_*ref*_ is the reference temperature (in ^*°*^*C*) at which the gating variable time constants were determined initially, and *T* is the operating temperature (in ^*°*^*C*) at which the experiment was conducted. The factor *Q*_10_ is an empirical measure that captures the temperature sensitivity of physiological processes and is defined as the fold change per 10^*°*^*C* from some reference temperature. It is experimentally determined by:

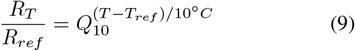

where *R*_*T*_, *R*_*ref*_ are the rates of processes at temperatures *T* and *T*_*ref*_ respectively. A temperature-independent process would therefore have a *Q*_10_ factor of 1. For reference, ion channel conductances are typically reported to have *Q*_10_ in the range of 1.2 to 1.5. Secondly, the voltage-dependent time constant for the gates, *τ*_*x*_(*V*_*m*_), is also expressed with temperature dependence. This dependency, too, is formulated as a *Q*_10_ factor which accounts for some of the observed variations. At higher temperatures, this correction factor reduces the time constant, effectively causing a quicker reaction to channel activation and inactivation. *Q*_10_ factors for ion channel activation or inactivation are typically in the range from 2.0 to 4.0.

The models that include temperature dependence may be based on experimental recordings of the channels; however, it is not guaranteed that the modelers have taken into account the difference in the *Q*_10_ factor between conductance and time constants. It is challenging to verify *post hoc*. Furthermore, the *Q*_10_ factor can also vary depending on the type of neuron and animal species. These nuances are also difficult to account for when ion channel models have been used interchangeably or with only minor changes. To test whether there was a temperature dependence included in the model, we ran the simulations for the original model at two different temperatures, as mentioned before, and asserted a temperature dependence if there was a non-zero area between the time constant curves at the two temperatures, or if there was a difference between the maximum conductance at the two temperatures. In the current database, 45% of the models exhibit some temperature dependence. 49% (1500 of 3083 tested) of all models show temperature dependence of the time constant, and 12% (358 of 3083 tested) of all models show temperature dependence for conductance. In many cases, the temperature dependence was entirely ignored; in these cases, we set the temperature fudge factor *Q*_10_ = 1.

### 4.5 Summary of the extracted gates

From all the voltage-dependent channels (3083 tested), we extracted 4190 gates (K:1629, Na:1338, Ca:806, Ih:170, KCa:247). Of these, 2233 were activating gates (steady-state value at −100 mV < steady-state value at 100 mV) and 1820 were inactivating; 137 gates could not be categorized because of other errors in extracting the gating variables.

### 4.6 Fitting omnimodels to extracted curves and computing errors

We used the Python library scipy ([37]) and *curve_fit* to fit the data points from the steady state and time constant curves to the omnimodels (Eqs. 1, 2, 3). The curve fitting algorithm was set to run up to 30000 iterations and gave us the coefficients of the omnimodel’s gates. To compare the errors between the raw data of the gates and our fits, we calculated three different errors. First of these was the root mean squared error (RMSE), computed as follows:

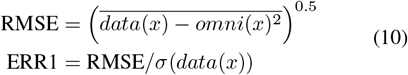

Here, *σ*(*data*(*x*)) corresponds to the standard deviation of the raw data (*data*) extracted at the membrane potentials *x* values. The corresponding values from the fitted curve are represented as *omni*(*x*). Secondly, we also computed the error based on the area between a cubic spline interpolation (*Spline*) of the raw data and the fitted curve (*omni*) as follows:

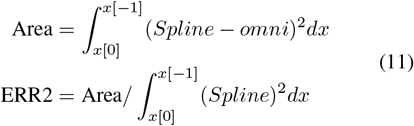

Where the notation *x*[0] and *x*[*−*1] indicate the lowest and highest membrane potentials at which the raw data were extracted. And finally, we also used Frobenius norm (indicated as ∥ ∥) of the difference between the data (*data*(*x*)) and the fitted curves (*omni*(*x*)), to compute an additional error metric as follows:

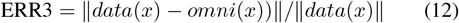

### 4.7 Creation of standardized .mod files and running ICG protocols and error evaluation

We developed a Python script that takes the coefficients of the omnimodels as a Python dictionary entry and produces a .mod file that can be run in the Neuron simulator. We could also choose to construct a simpler omnimodel which uses its steady state curve based on Eq.1 instead of 2, with the time constant curve for both models set to the third order (Eq.3, with *e*1, *e*2 set to 0). We used this newly generated omnimodel mod file and repeated the ICG protocols, which were first presented in our earlier work [8]. The resulting currents were compared with the original model output. For the comparison, we used the same three error methods from before (Eqs. 10, 11, 12). There were no significant differences in outcome between the three error methods that we used.

### 4.8 Details of single-neuron simulations

We tested two previously published pyramidal neuron models by systematically replacing their original ion channel mechanisms with corresponding omnimodels, when available. For the layer 5 pyramidal neuron model [10], we substituted 8 of the 10 ion channels with omnimodels and validated the implementation using three published stimulation protocols. For the layer 2/3 pyramidal neuron model [11], we performed similar substitutions and examined neuronal responses to depolarizing current pulses and varying levels of somatic depolarization.

### 4.9 ICG explorer and visualizer

In addition to the extraction of gating variables, we evaluated the chronological lineage and prevalence of channel mechanism reuse among ion channels curated in ModelDB [7, 38]. This approach was motivated by the observation, confirmed by inspection of fitted omnimodel coefficients, that gating variables form tight clusters, consistent with the extensive reuse of near-identical (and often identical) channel mechanisms.

To establish a practical pipeline for prioritizing channel candidates for modeling and fitting, we developed a source-code similarity analysis that produces a canonical, compilation-equivalent string for each NMODL file. This normalization process involves three steps: 1. Stripping comments (including those inside `VERBATIM` blocks), 2. Removing non-significant whitespace, and 3. flattening trivial formatting. We then computed the pairwise Levenshtein similarity [39] on these canonical strings. Files that were byte-identical were merged into “clone pools” of functionally identical implementations. Pools are connected to their nearest neighbors to infer possible code reuse whenever the similarity ratio exceeds a user-defined cutoff.

To make this analysis broadly accessible, we provide an interactive web application (https://icg-explorer.org/visualizer) that renders the similarity graph. This visualization utilizes a D3.js force-directed layout [40], which organizes channels into clusters based on their textual similarity, with clusters mutually repelled by a gravity-force algorithm. Within each byte-identical clone pool, we mark a canonical model as the earliest publication (identified by the lowest ModelDB ID) to represent the chronological parent. Graph connections are inferred from the pairwise string similarity and are thus tentative; they suggest plausible development paths through successive minor edits rather than asserting true ancestry. The interface supports pairwise and multi-selection side-by-side diffs (keyboard shortcut: d), rendering HTML tables with color-coded changes (additions: green, deletions: red, substitutions: yellow). These features allow users to: a) identify clusters of similar source-code nodes, b) trace plausible development paths of the ion channel models, and c) prioritize work by keeping one canonical model per clone pool, reviewing near-duplicates first, and skipping re-fitting or re-validation on functionally identical code.

## Acknowledgements

We thank Andrea Navas-Olive and Douglas Feitosa Tomé for their feedback on the manuscript.

## Author contributions

CC, WP, performed all the simulations, and PB created the ion channel explorer. PJG, J-ML, and JHM conceptualized omnimodels. CC, WP, and PB generated the figures. The manuscript was written by CC, WP, PB, and edited by CC, WP, PB, PJG, J-M.L, JHM, and TPV.

## Data and code availability

Data and code is available at the ICG Github repository and new features at ICG Explorer.

